# Readiness of nociceptor cell bodies to generate spontaneous activity results from background activity of diverse ion channels and high input resistance

**DOI:** 10.1101/2023.06.30.547260

**Authors:** Jinbin Tian, Alexis G. Bavencoffe, Michael X. Zhu, Edgar T. Walters

## Abstract

Nociceptor cell bodies generate “spontaneous” discharge that can promote ongoing pain in persistent pain conditions. Little is known about the underlying mechanisms. Recordings from nociceptor cell bodies (somata) dissociated from rodent and human dorsal root ganglia (DRGs) have shown that prior pain in vivo is associated with low-frequency discharge controlled by irregular depolarizing spontaneous fluctuations of membrane potential (DSFs), likely produced by transient inward currents across the somal input resistance. Here we show that DSFs are associated with high somal input resistance over a wide range of membrane potentials, including depolarized levels where DSFs approach action potential (AP) threshold. Input resistance and both the amplitude and frequency of DSFs were increased in neurons exhibiting spontaneous activity. Ion substitution experiments indicated that the depolarizing phase of DSFs is generated by spontaneous opening of channels permeable to Na^+^ and/or Ca^2+^, and that Ca^2+^-permeable channels are especially important for larger DSFs. Partial reduction of the amplitude and/or frequency of DSFs by perfusion of pharmacological inhibitors indicated small but significant contributions from Nav1.7, Nav1.8, TRPV1, TRPA1, TRPM4, and N-type Ca^2+^ channels. Less specific blockers suggested a contribution from NALCN channels, and global knockout suggested a role for Nav1.9. The combination of high somal input resistance plus background activity of diverse ion channels permeable to Na^+^ and/or Ca^2+^ produces DSFs that are poised to reach AP threshold if resting membrane potential (RMP) depolarizes, AP threshold decreases, and/or DSFs become enhanced -- all of which have been reported under painful neuropathic and inflammatory conditions.

## 1. Introduction

Primary nociceptors and low-threshold mechanoreceptors are normally electrically silent, but in neuropathic pain conditions action potentials (APs) may be generated “spontaneously” to drive ongoing pain in rodents and humans ^16, 48, 65, 68, 69^. Electrophysiological studies have shown a link between suprathreshold spontaneous activity (SA) in dorsal root ganglion (DRG) neurons and spontaneous fluctuations of resting membrane potential (RMP). Recordings using sharp electrodes from the large cell bodies (somata) of A-fibers within excised DRGs revealed high-frequency, sinusoidal oscillations that could generate regular tonic or bursting discharge ^3, 71^. However, sharp-electrode recordings from the somata of C-fibers reveal much smaller spontaneous subthreshold membrane potential fluctuations which fail to reach AP threshold because shunting around the impaling electrode dramatically reduces input resistance in smaller cells – an effect that has probably led to an underestimate of how common somally generated SA is in nociceptors. Whole-cell patch recording (with gigaohm membrane seals) from somata of dissociated C-fiber neurons reveals much larger spontaneous fluctuations of RMP that may reach AP threshold in painful conditions ^51, 61^. These fluctuations differ from the sinusoidal oscillations in A-fiber somata in their lower frequency and irregular properties. Amplitudes both of the sinusoidal oscillations in large DRG neurons ^3, 71^ and the irregular depolarizing spontaneous fluctuations (DSFs) in small DRG neurons ^9, 42, 51^ increase in neuropathic pain models. Importantly, in human patients with neuropathy associated with cancer, irregular DSFs and the incidence of dissociated neurons exhibiting SA were selectively increased in cells taken from excised DRGs corresponding to painful dermatomes ^49, 50^.

Sinusoidal oscillations in large DRG neurons can be eliminated by substituting extracellular NaCl with choline-Cl and by concentrations of tetrodotoxin (TTX) that selectively block TTX-sensitive voltage-gated sodium channels (VGSCs) ^3, 71^. Blockers of voltage-gated calcium channels (VGCCs) and voltage-gated potassium channels had no effect on the oscillations ^2, 3^. This suggested a mechanism in which subthreshold sinusoidal oscillations are generated by reciprocating actions of depolarizing currents through TTX-sensitive VGSCs and steady repolarizing current through potassium leak channels ^39, 40^. In contrast, mechanisms underlying irregular DSF generation have not been reported for any neuron type ^65^.

Because of the prominence of DSFs in nociceptors and their close association with multiple forms of neuropathic pain in rodents and humans ^6, 9, 42, 50, 51^, we have investigated the contributions of passive membrane properties and of different ions and ion channels to the generation of DSFs in nociceptors dissociated from normal, uninjured mice. While much is known about the contributions of numerous ion channels to RMP in rodent nociceptors ^18^, the ion channels important for DSF generation have not been identified. Our findings reveal distinctive electrophysiological properties that enable silent nociceptors to readily transition to continuing, irregular electrical activity. Spontaneous background activity of diverse ion channels permeable to Na^+^ and/or Ca^2+^ combined with high somal input resistance produces irregular DSFs that are poised to drive ongoing discharge under painful conditions that increase nociceptor excitability by sustained depolarization, increased activity of any of these channels, and/or additional increases in input resistance.

## 2. Methods

### 2.1. Animals

All procedures followed the guidelines of the International Association for the Study of Pain and were approved by the McGovern Medical School at UTHealth Animal Care and Use Committee. Totals of 90 male and 30 female wild-type C57BL/6 mice were included in this study. Although male and female mice were included in all experiments, this study was not designed to investigate sex differences. Thus, only large sex differences would be detected with the sample sizes used, and data from both sexes were always pooled. Weak suggestions of sex differences occurred in some of the results, and these are noted in the text. Adult mice (16-30 g, 1-5 per cage) were allowed to acclimate inside the McGovern Medical School animal research facility for at least 4 days before beginning experiments. Mice were provided with food and water ad libitum. Male and female Nav1.9^+/−^ mice were purchased from Jackson Laboratory (B6.129P2-Scn11atm1Dgen/J Strain #: 005837). Tissue samples of offspring mice were subjected to a standard PCR genotyping assay (Jackson Laboratory genotyping protocol 23186). Four male and 4 female Nav1.9^−/−^ mice were used in this study.

### 2.2. Dissociation and culture of primary sensory neurons

An in vitro preparation was used to investigate DSFs and other electrophysiological properties of dissociated mouse DRG neurons, using methods previously described ^9, 42^. Mice were euthanized with isoflurane and intracardially perfused with ice-cold PBS. Dorsal root ganglia were quickly and carefully removed and trimmed in ice-cold DMEM, and digested with trypsin (0.3 mg/ mL, Worthington Biochemical Corporation, Lakewood, NJ, #LS003702) and collagenase D (1.5 mg/mL, Sigma, #11088858001) for 30 minutes at 34 °C. Debris was removed by 2 successive centrifugations (6 minutes at 600 rpm) and cells were plated onto coverslips coated with 0.01% poly-L-ornithine (Sigma, P4957) and incubated overnight at 37 °C with 5% CO_2_ in serum-free DMEM.

### 2.3. Recording from dissociated DRG neurons

Whole-cell patch clamp recordings were performed at room temperature 18–30 hours after dissociation using an EPC10 USB (HEKA Elektronik, Lambrecht/Pfalz, Germany) amplifier. Patch pipettes were made of borosilicate glass capillaries (Sutter Instrument Co., Novato, CA) using a Narishige PC-10 puller, and fire-polished with an MF-830 microforge (Narishige, Tokyo, Japan) to a final pipette resistance of 3–8 MΩ when filled with an intracellular solution composed of (in mM): 134 KCl, 1.6 MgCl_2_, 13.2 NaCl, 3 EGTA, 9 HEPES, 4 Mg-ATP, and 0.3 Na-GTP, which was adjusted to pH 7.2 with KOH and to 300 mOsm. Isolated small neurons with a soma diameter ≤30 µm were observed at 40x magnification on an IX-71 (Olympus, Tokyo, Japan) inverted microscope and recorded in an extracellular solution (ECS) containing (in mM): 140 NaCl, 3 KCl, 1.8 CaCl_2_, 2 MgCl_2_, 10 HEPES, and 10 glucose, which was adjusted to pH 7.4 with NaOH and to 320 mOsm. In ion substitution experiments, the driving force for Na^+^ and Ca^2+^ currents at a holding potential of −45 mV was greatly reduced by lowering extracellular Na^+^ concentration to 2.2 mM (low-Na^+^ ECS), and combining 0 mM (nominally) extracellular Ca^2+^ with 1 mM EGTA (low-Ca^2+^ ECS). To maintain osmolarity and minimize charge screening effects, choline-Cl replaced most of the NaCl in low-Na^+^ ECS, and 1.8 mM MgCl_2_ replaced Ca^2+^ in low-Ca^2+^ ECS. Low-Na^+^ ECS contained (in mM) 2.2 NaCl, 137.8 choline-Cl, 3 KCl, 1.8 CaCl_2_, 2 MgCl_2_, 10 HEPES, and 10 glucose. Low-Ca^2+^ ECS contained 140 NaCl, 3 KCl, 3.8 MgCl_2_, 1 EGTA, 10 HEPES, and 10 glucose. Combined low-Na^+^, low-Ca^2+^ ECS contained 2.2 NaCl, 137.8 choline-Cl, 3 KCl, 3.8 MgCl_2_, 1 EGTA, 10 HEPES, and 10 glucose.

Whole-cell patch clamp recording commenced after obtaining a tight seal (> 3 GΩ) and rupturing the plasma membrane under voltage clamp at −60 mV. Recordings were acquired with PatchMaster v2x73 (HEKA Elektronik, Lambrecht/Pfalz, Germany). The liquid junction potential was calculated to be ~4.3 mV and not corrected, meaning the actual potentials may have been ~4.3 mV more negative than indicated in the recordings and measurements presented herein.

### 2.4. Quantifying depolarizing spontaneous fluctuations of membrane potential (DSFs)

DSFs were analyzed as previously described ^51^. Briefly, we used a custom program (SFA_pub.py) to quantify irregular DSFs in patch recordings, imported as time and voltage coordinate data for 30-40 second periods from recordings obtained with PatchMaster sampled at 10-20 kHz and filtered with a 2.9 kHz-Bessel filter. The program used a sliding median function to calculate RMP at each point and return coordinates, amplitudes, and durations of identified APs and DSFs (minimum amplitude 1.5 mV, minimum duration 5 ms) and hyperpolarizing spontaneous fluctuations of membrane potential (HSFs), as well as a continuous color-coded plot of membrane potential generated using the matplotlib library (Python v3.6, Python Software Foundation, Beaverton, OR) ^51^. Manual inspection of each plot confirmed that each AP was generated by a suprathreshold DSF. Conservative estimates of the amplitudes of suprathreshold DSFs were measured as the most depolarized potential reached by the largest subthreshold DSF recorded at the indicated holding potential ^51^. During DSF analysis, APs were substituted by the largest subthreshold DSF measured within the same 30-40 second recording period. Recordings with higher frequency firing rates (>0.5 Hz over the entire period) were excluded because of interactions of DSFs with AP afterhyperpolarizations and because fewer large subthreshold DSFs are available for measurement when most of the larger DSFs are suprathreshold.

### 2.5. Low-frequency voltage clamp

This method was used to prevent slow changes in membrane potential without attenuating DSFs. Low-frequency voltage clamp (PatchMaster v2x73, HEKA Elektronik) acts as a current clamp for faster signals and a voltage clamp for slower signals, using a low-pass filtered recording of membrane potential as a constant averaged command voltage to control feedback injection of current to maintain the set holding potential. We chose an effective feedback speed of ~3 seconds, which adequately clamped the holding potential without producing detectable effects on DSFs or APs. In these studies, we clamped the holding potential to ~-50 mV or ~-45 mV.

### 2.6. Pharmacological agents

Ion channel blockers, including ProTX II, ononetin, CBA and ZD-7288 were purchased from Tocris Bioscience; gadolinium chloride, L-703606, and capsaicin from Sigma-Aldrich; TTA-P2 and ω-conotoxin GVIA from Alomone Labs; A-967079 and Pico-145 from MedChemExpress; A-803467 from Apexbio Technology LLC; AMG-9810 from Cayman Chemical Company; nimodipine from Ascent Scientific Ltd.; choline-Cl from MP Biomedicals; EGTA from Fisher Scientific. Drugs were applied through a gravity-driven multi-outlet device with the selected outlet placed ~50 μm away from the cell being recorded. Drugs were diluted to the final concentration in ECS and applied at approximately 0.25 ml/min in a recording chamber volume of 3.5 ml. All recordings were made at room temperature (22-23°C).

### 2.7. Data analysis

Statistical analyses of raw electrophysiological data and SFA_pub.py output were performed using Prism V9.5.1 (GraphPad Software, La Jolla, CA). Averages are presented as mean or median values. All datasets were tested for normality with the D’Agostino Pearson omnibus normality test. Normally distributed data were tested with parametric tests: either paired or unpaired *t* tests, or 1-way ANOVA for repeated measures followed by Tukey’s multiple comparisons test. In some cases, Bonferroni corrections were made for multiple comparisons. Single comparisons of data that were not normally distributed were tested with a Mann-Whitney U test. Statistical significance was set at *P* < 0.05, and all reported values are 2-tailed. Specific statistical tests corresponding to each figure are listed in the figure and table legends.

## 3. Results

### 3.1. Dissociated mouse DRG neurons exhibit high input resistance across a wide range of membrane potentials

By Ohm’s Law, the amplitudes of fluctuations in membrane potential occurring spontaneously in dissociated nociceptor somata under whole-cell current clamp are determined by changes in the product of the net current flowing across the plasma membrane and the neuron’s input resistance. Higher input resistance will increase changes in membrane potential caused by spontaneous opening and closing of ion channels in the soma membrane. We measured input resistance in nonaccommodating (NA) DRG neurons ^51^ (the vast majority of neurons sampled) under whole-cell current clamp at holding potentials of approximately −70, −60, and −50 mV, spanning much of the range of RMP in dissociated mouse nociceptors ^9, 42^. Conventional measurement of input resistance using a series of increasingly negative 500-ms constant-current pulses (−5 pA increments) evoked hyperpolarizations that were similar for the same currents injected from each holding potential (**Fig. 1A**) and showed little voltage dependence (**Fig. 1B**). The hyperpolarizing voltage changes corresponded in most cases to an input resistance of 3 to 4 GΩ, which is quite high compared with values reported for most neurons during whole-cell patch recording (see Discussion). As expected, input resistance showed a negative correlation with input capacitance (**Fig. 1C**), which is a measure of total membrane area and is correlated with soma diameter.

**Figure 1.**
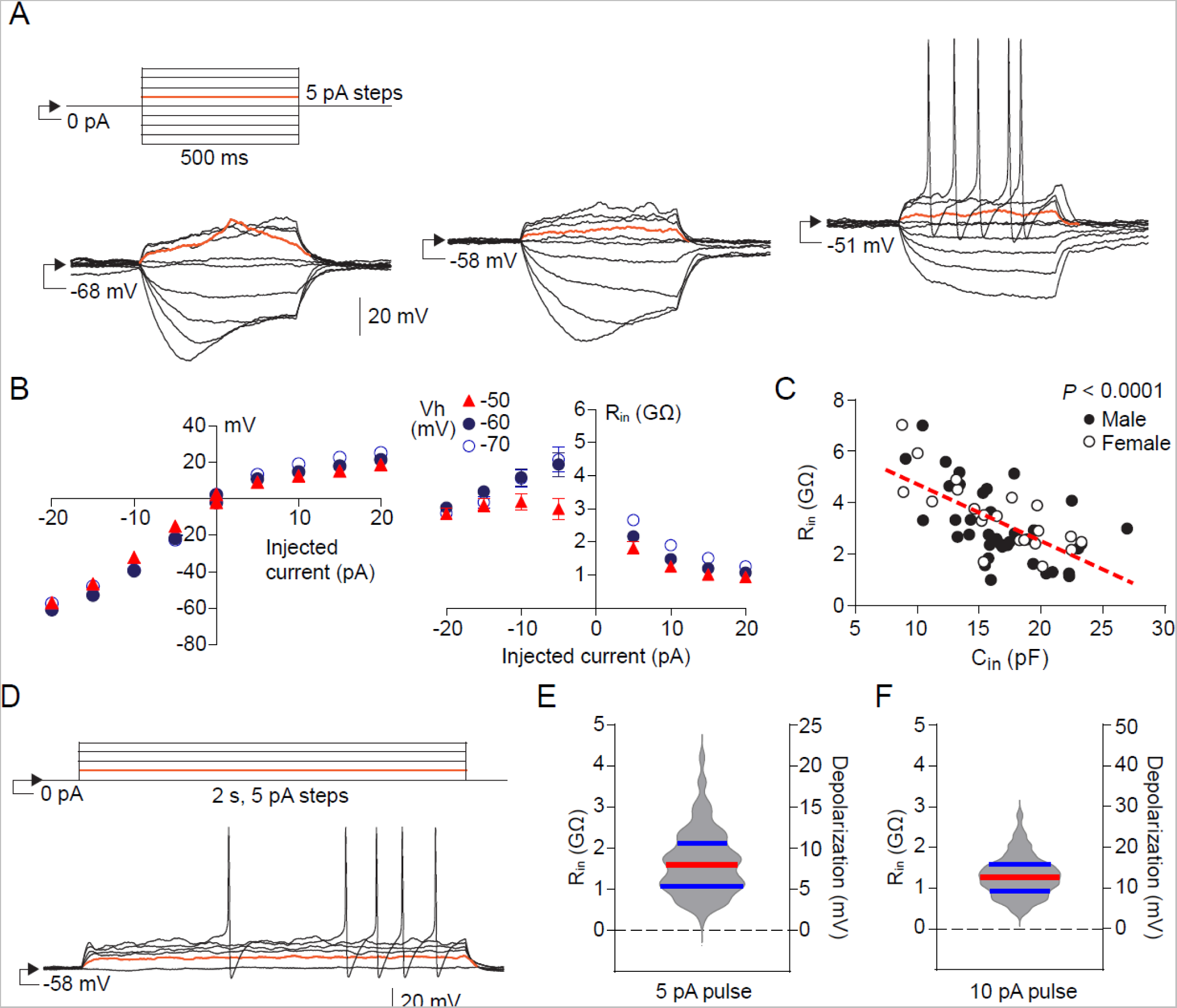
Dissociated mouse DRG neuron somata exhibit high input resistance across a wide range of holding potentials. (A) Representative examples of voltage responses to constant current pulses (5-pA increments) of negative and positive current delivered under whole-cell current clamp from holding potentials of approximately −70, −60, and −50 mV. Increasingly negative steps were delivered first, followed by increasingly positive steps, and each sequence began with a 0-pA sweep. Input resistance (R_in_) was calculated from the minimum hyperpolarizing potential (excluding sag), or the plateau value in each depolarizing potential. The response to the positive 5-pA step in each set of traces is colored red. The 5-pA response on the left shows an apparent DSF superimposed on the depolarizing plateau. Estimates of the plateau values used to calculate R_in_ excluded any obvious DSFs. (B) The left panel shows the mean voltage responses for all neurons tested (n = 37 for each holding potential), revealing a relatively linear voltage-current (V-I) relationship for hyperpolarizing pulses and smaller, non-linear voltage responses for depolarizing pulses. The right panel plots the mean R_in_ calculated for each current pulse and holding potential. (C) Inverse correlation between Rin and input capacitance (C_in_) for the neurons in parts A and B (n = 59). Pearson correlation, r = −0.6355, *P* < 0.001. (D) Example of voltage responses to positive constant current pulses used to calculate R_in_ and determine rheobase. The calculated R_in_ and corresponding depolarization for each neuron tested are shown for the 5-pA pulse (n = 148) (E) and 10 pA pulse (n = 200) (F). The median value for all neurons in each case is indicated by the red line, and interquartile range by the blue lines. R_in_, input resistance; V_h_, holding potential.

DSFs are significant for pain because they can bridge the gap between RMP and AP threshold to drive ongoing discharge in nociceptors. Thus, the effective input resistance between RMP and AP threshold, i.e., as measured with depolarizing pulses, is most important for DSF function. As shown in **Figures 1A** and **1B**, positive current pulses produced depolarizations that exhibited a nonlinear voltage dependence, as would be predicted by the expression in DRG neurons of numerous voltage-gated ion channels that open in response to depolarization, thereby decreasing input resistance but potentially triggering active responses that lead to AP generation. Interestingly, the smallest applied depolarizing pulse, 5 pA, rarely evoked APs (**Fig. 1A**) while producing a steady-state depolarization of ~10 mV from each of the tested holding potentials (**Fig. 1B**), representing an input resistance of ~2 GΩ in this voltage range. A 10-mV depolarization would correspond to a very large DSF amplitude in mouse DRG neurons ^9, 42^. To expand the number of neurons sampled for input resistance analysis in the depolarized subthreshold voltage range, we made use of the test pulses serving to measure rheobase and to distinguish NA neurons from rapidly accommodating (RA) neurons ^51^, which included 2-second depolarizing steps of 5 and 10 pA (**Fig. 1D**). These pulses produced median steady-state depolarizations of 8.0 and 12.6 mV, respectively, corresponding to median input resistances of 1.6 and 1.3 GΩ (**Fig. 1E, 1F**), confirming the DRG neurons’ high input resistance in the voltage range where large DSFs can sometimes reach AP threshold. No significant differences were found in the input resistances of neurons taken from male mice (median 2.7 GΩ, n = 39) and female mice (3.5 GΩ, n = 20) using hyperpolarizing pulses (**Fig. 1C**), or with depolarizing pulses (e.g., for the 10-pA pulse, males, 1.3 GΩ, n = 114; females, 1.2 GΩ, n = 86, **Fig. 1F**), Mann-Whitney U test in each case.

### 3.2. DSFs in dissociated mouse DRG neurons exhibit irregular waveform properties

As previously shown in mice, dissociated small and medium-sized DRG neurons exhibit irregular, spontaneous, subthreshold fluctuations of membrane potential across a wide range of RMPs (−40 to −80 mV), which increase in amplitude under neuropathic conditions (spinal cord injury and chemotherapy models) ^9, 42^. However, the waveforms, frequencies, durations, and patterns of occurrence of DSFs were not described. Representative waveforms observed in raw recordings from neurons with smaller and larger spontaneous fluctuations of membrane potential, respectively, are illustrated in **Figures 2A** and **B**. Peak-to-peak amplitude variation was 2.0 mV and 8.2 mV in these two samples. Also evident are the large ranges of 1) positive and negative peak values, 2) intervals between peaks, and 3) waveform durations, as well as apparent summation of waveforms. We then recorded from a non-excitable cell type, cultured primary human umbilical vein endothelial cells (HUVECs), exhibiting high input resistance (~2 GΩ) ^63^ similar to what we found in mouse DRG neurons. The high input resistance and depolarized RMP in HUVECs reflect the very low resting current through available K^+^ channels and paucity of other open channels ^63^. We found slow peak-to-peak fluctuations of ~1.5 mV in HUVECs (**Fig. 2C**), which are smaller than most of the peak-to-peak fluctuations recorded from mouse NA neurons under identical conditions (e.g., **Fig. 2A, B**, and see below). The fluctuations observed in HUVECs and DRG neurons are almost devoid of typical signs of electrical noise, including very high frequency spikes or 60 Hz hum, indicating that much of the fluctuating signal is biological. Furthermore, when extracellular Na^+^ and Ca^2+^ were reduced (see section 3.4), peak-to-peak fluctuations often were < 1 mV, indicating that an analytic cutoff ≥ 1 mV peak-to-peak would safely avoid electrical noise components to reveal biological fluctuations.

**Figure 2.**
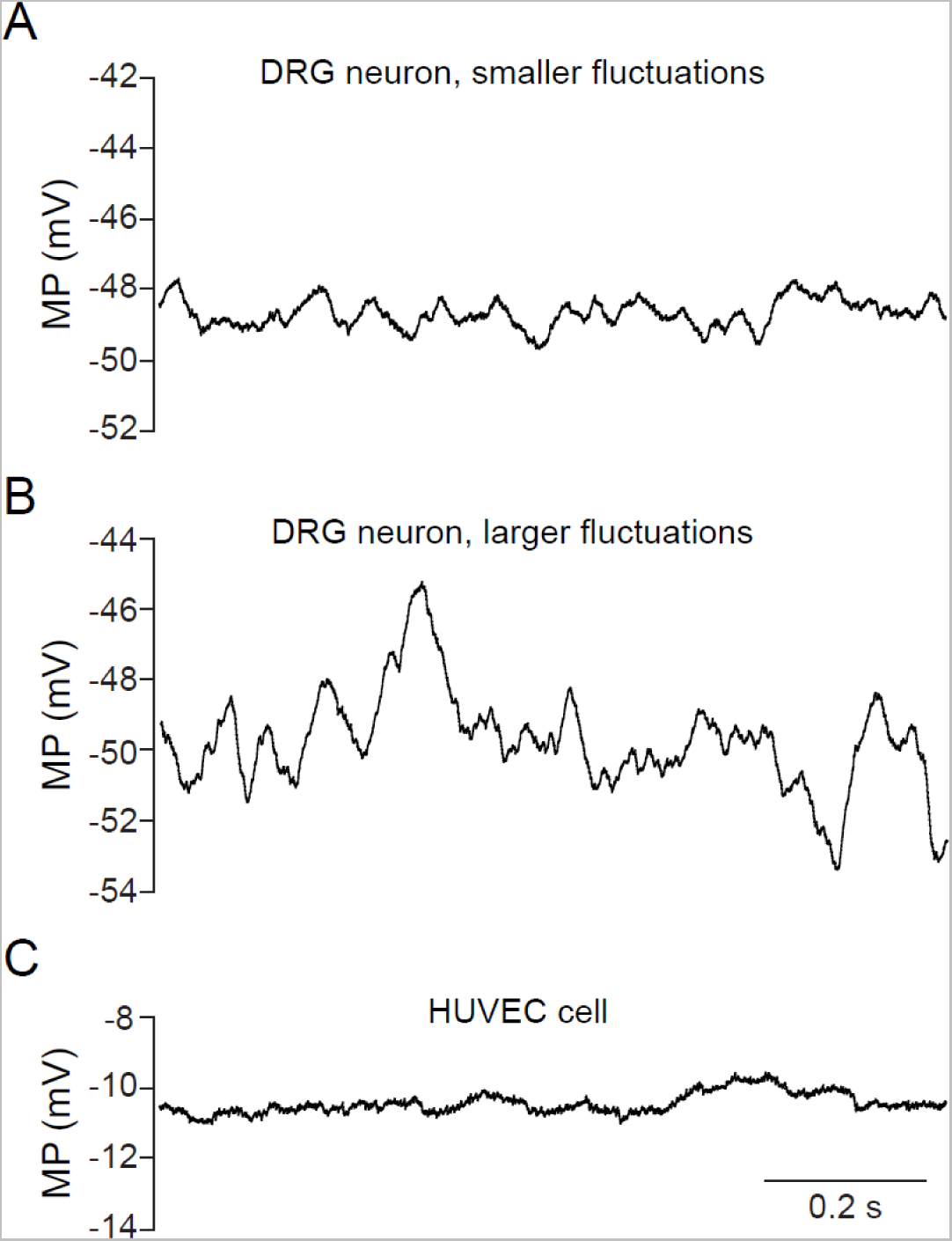
Irregular properties of spontaneous fluctuations of RMP in dissociated mouse DRG neuron somata. (A) Example of a neuron with smaller fluctuations. Note the irregular waveforms, amplitudes, and inter-peak intervals. (B) Example of a neuron with larger fluctuations that also are highly irregular. (C) Example of spontaneous fluctuations in a non-excitable HUVEC cell having an input resistance similar to that of smaller mouse DRG neurons. DRG, dorsal root ganglion; DSF, depolarizing spontaneous fluctuation; HUVEC, human umbilical vein endothelial cell; RMP, resting membrane potential.

To quantify DSF properties, we used an automated algorithm to selectively measure the depolarizing fluctuations relative to a sliding median computed for RMP ^51^, as shown in **Figure 3A**. As done previously for mouse, rat, and human DSFs, we set a minimum cutoff of 1.5 mV deviation from the sliding median (corresponding to ~3 mV peak-to-peak), which is well above the electrical noise level and above nearly all the peak-to-peak fluctuations observed in HUVECs (**Fig. 2C**). DSFs accepted for analysis are colored red. Larger DSFs were defined as those >3 mV (green dashed line). Also indicated are hyperpolarizing spontaneous fluctuations (HSFs, colored blue), which were not analyzed in this study. The ranges and median values of the amplitudes, frequencies, and durations of DSFs recorded at RMP under current clamp are shown in **Figures 3B, C**, and **D**. A small sex difference was suggested by significantly lower median DSF amplitudes in samples from females (1.9 mV, n = 29) versus males (2.1 mV, n = 2.1 mV, *P* = 0.01) but this difference may be explained by a larger median input capacitance in the female than male samples (19.8 versus 15.7 pF, *P* = 0.02) (Mann-Whitney U test in each case). In the neurons taken from previously uninjured mice, DSF amplitude was not significantly correlated with RMP (**Fig. 3E**, see also ^9, 51^), while DSF frequency was positively correlated (**Fig. 3F**) and DSF duration was negatively correlated with RMP (**Fig. 3G**).

**Figure 3.**
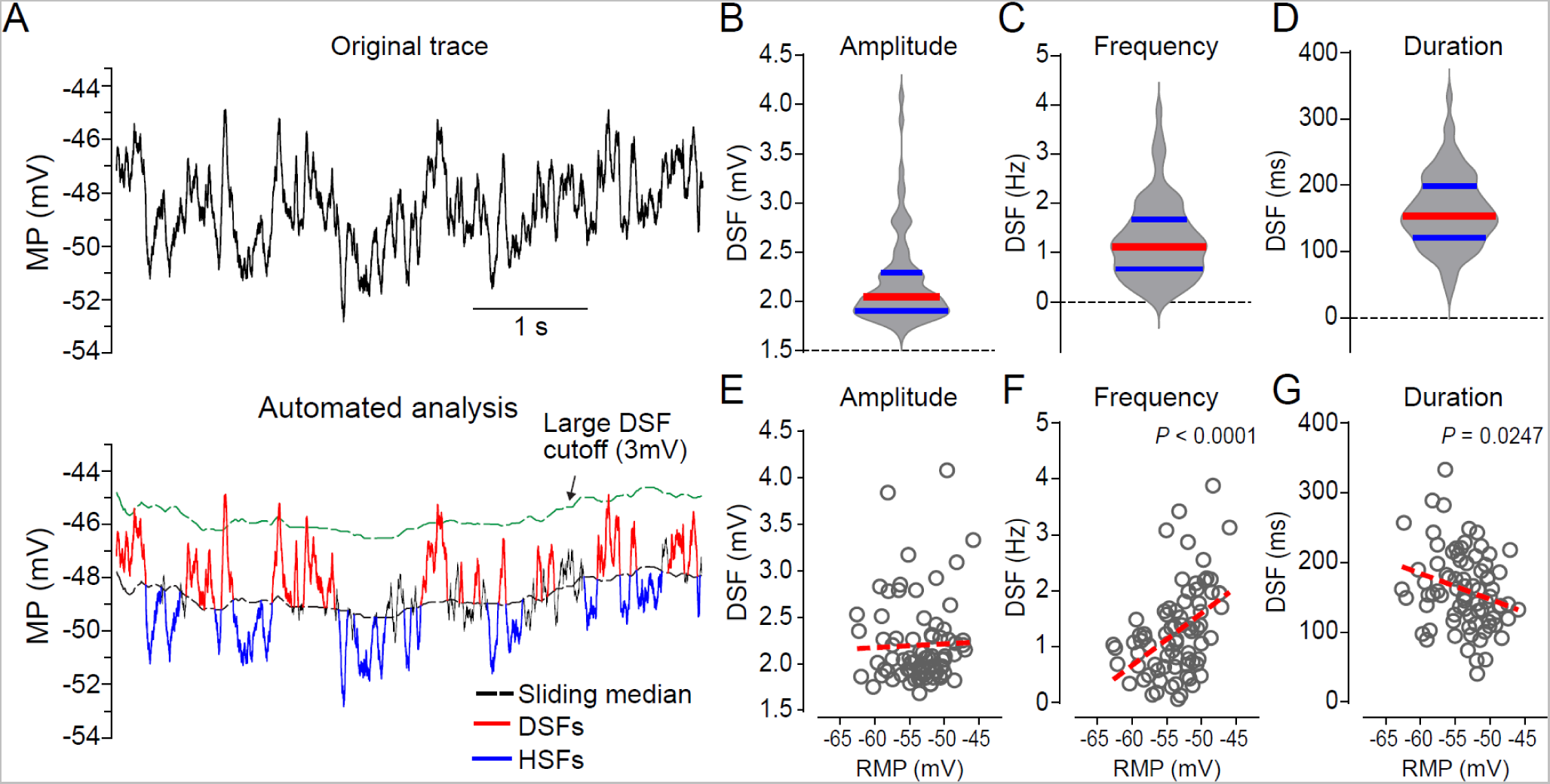
Basic properties of DSFs recorded in DRG neurons dissociated from previously uninjured mice. (A) Automated measurement of DSFs. The original recording is shown above the output of the program used to compute the sliding median of the fluctuating RMP (black line) and to measure depolarizing fluctuations (DSFs) and hyperpolarizing fluctuations (HSFs) having minimum absolute amplitudes of 1.5 mV (red and blue segments, respectively). The green line shows the running 3 mV value used to define larger DSFs. Measured fluctuation properties revealed considerable variation across the neurons tested (n = 79) in DSF amplitude (B), DSF frequency (C), and DSF duration (D). Red lines indicate medians, blue lines interquartile range. Each data point composing the violin plot represented the mean value for all DSFs recorded from 1 neuron across 30-40 seconds. Relationships to the RMP of each recorded neuron are shown for DSF amplitude (E), DSF frequency (F), and DSF duration (G). Red dashed lines show linear regression. *P* values were derived from Pearson correlation, r = 0.4439 (E) and r = −0.2426 (F). DSF, depolarizing spontaneous fluctuation; MP, membrane potential; RMP, resting membrane potential.

### 3.3. Input resistance and DSF amplitude are increased in DRG neurons exhibiting spontaneous activity (SA)

In our studies of spinal cord injury (SCI), we found the incidence of dissociated rodent DRG neurons exhibiting SA to be greatly increased by SCI (usually exceeding 50%), but a lower incidence of SA (usually <20%) sometimes occurs in neurons from previously uninjured, naïve animals ^5, 7, 9, 51^. Interestingly, the electrophysiological properties of SA neurons examined previously from naïve and SCI rats did not differ, suggesting that while SCI potently induces a hyperactive state that can drive SA at RMP in nociceptors, the same state is occasionally induced in neurons from uninjured animals ^7^, perhaps in response to injury signals produced during DRG excision and neuronal dissociation ^75^. Thus, we asked how electrophysiological properties of DRG neurons from previously uninjured mice might differ in neurons with and without SA. Classification of the neurons into the SA or No SA (silent) groups was based on whether any spontaneous APs occurred at the neuron’s RMP during the early 60-second monitoring period prior to subsequent tests. Indeed, we found some mouse DRG neurons had SA resembling the SA previously examined after SCI ^9^ (**Fig. 4A**). These SA neurons in previously uninjured mice had significantly depolarized RMP compared to neurons without SA (**Fig. 4B**), their input resistance was significantly greater (**Fig. 4C**), and their rheobase (the current needed to reach AP threshold during prolonged depolarization) was significantly lower (**Fig. 4D**), all of which are likely to contribute to the hyperactivity of these neurons. In the No SA group but not the SA group, neurons from female mice (n = 58) had significantly lower input resistance when tested with 10 pA pulses (*P* = 0.005, Mann-Whitney U test) than neurons from male mice (n = 70). This suggests a possible sex difference in nociceptor input resistance, although at least part of the difference in these samples might have resulted from larger input capacitance in the female than male groups (medians, 17.5 pF versus 16.0 pF).

**Figure 4.**
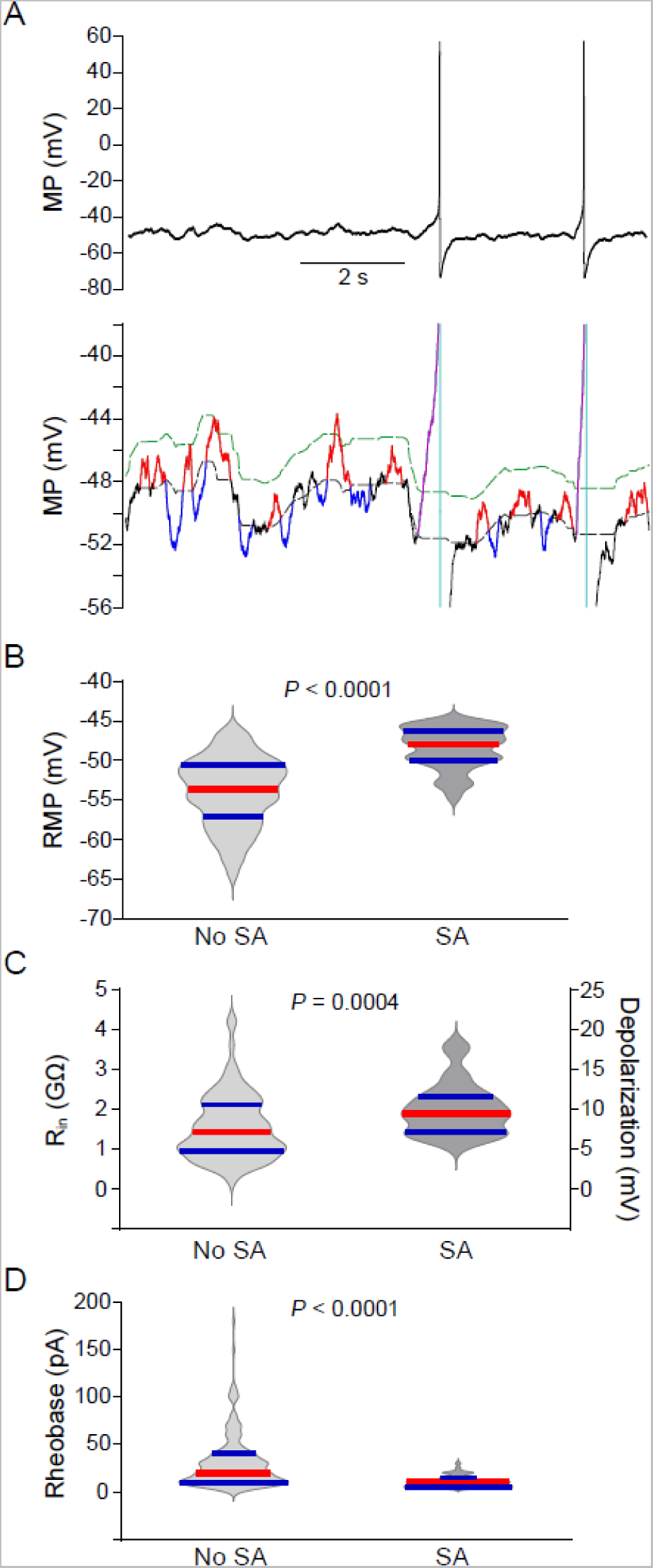
DRG neurons from previously uninjured mice can exhibit SA similar to that reported in neuropathic pain conditions. (A) Example of SA neuron and its measured DSFs. The color scheme is the same as in Figure 3A. (B) Significant depolarization of RMP of SA neurons (n = 58) compared with neurons lacking SA (“No SA” group, n = 178). (C) Significant enhancement of input resistance in SA neurons (n = 37) versus No SA (n = 91) neurons. (D) Significant reduction of rheobase in SA neurons (n = 56) versus No SA neurons (n = 176). Red lines indicate group medians, blues lines interquartile range. Each data point composing the violin plot represented the mean value for all DSFs recorded from 1 neuron across 30-40 seconds. Each group comparison was made using a Mann Whitney U test. MP, membrane potential; R_in_, input resistance; SA, spontaneous activity.

Because of the correlation between SA incidence and DSF amplitude found in rat and human DRG neurons ^50, 51^, we predicted that DSFs in mouse DRG neurons would be enhanced in SA neurons held at membrane potentials where APs are rarely generated but where the largest DSFs can sometimes reach AP threshold. We found previously that the incidence of neurons exhibiting AP generation (ongoing activity, OA) during prolonged artificial depolarization under current clamp to approximately −45 mV (as might occur during inflammation in vivo ^67^) provides a highly sensitive and physiologically relevant measure of hyperexcitability in mouse and rat DRG neurons ^9, 44^. Thus, we measured DSFs in neurons held at potentials of either −45 or −50 mV for 40 seconds. Holding potentials of −50 mV were used when any discharge occurring at a holding potential of −45 mV was of sufficient frequency (≥ 2 Hz) to interfere with measurement of DSFs. A few neurons that had RMPs positive to −45 mV were excluded so that none of the tested neurons had to be artificially hyperpolarized from their RMP to the test holding potential. Because some neurons depolarized slowly during prolonged current clamp, and both SA incidence and large DSF frequency increase at depolarized RMPs ^51^, in the these and the following experiments on DSFs we stabilized holding potentials for long periods (up to several minutes) by using low-frequency voltage clamp to prevent slow changes in membrane potential while minimally affecting DSFs and APs (**Fig. 5A**). We tested DSFs at holding potentials of −45 or −50 mV, but classification of the neurons into the SA or No SA group was based on whether any spontaneous APs occurred at the neuron’s RMP prior to the other electrophysiological tests. We found that DSFs were significantly larger in SA neurons than in No SA neurons during 40-second recordings at potentials clamped to −45 or −50 mV for both ranges of DSF amplitudes (1.5 – 3 mV and > 3 mV) (**Fig. 5B**). In addition, the frequency of larger DSFs (> 3 mV) was significantly higher in SA than No SA neurons (**Fig. 5C**).

**Figure 5.**
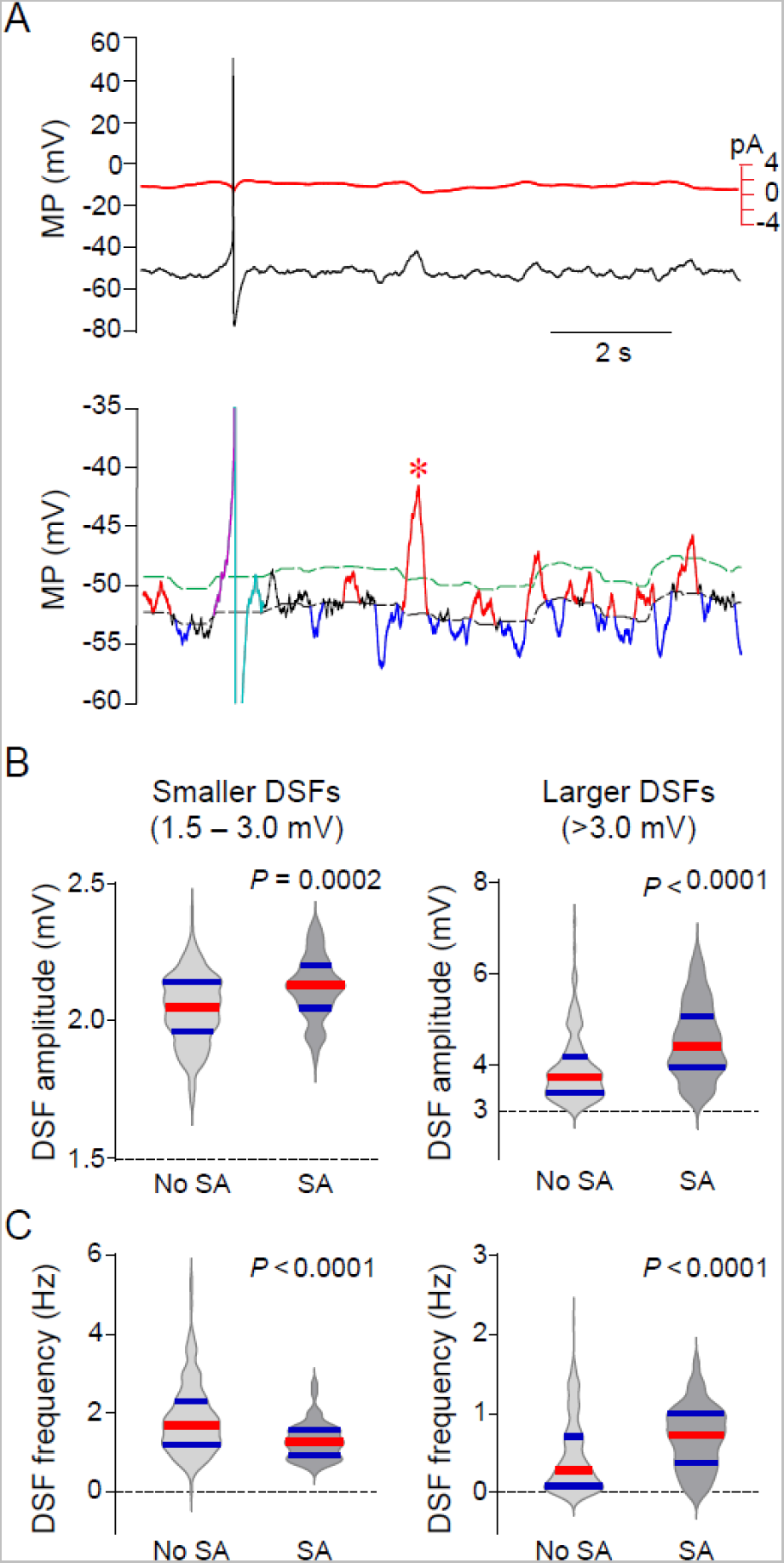
Hyperactive DRG neurons dissociated from previously uninjured mice have larger DSFs than electrically silent neurons from the same mice. (A) Example of a neuron exhibiting ongoing AP discharge (spontaneous activity, SA, a measure of hyperactivity) when held under low-frequency voltage clamp at ~-50 mV. The red line in the top panel indicates the changes in membrane current (negative is inward) associated with the imposed slow clamp of holding potential. The asterisk in the lower panel shows the largest DSF measured in this 10-second example. Because APs are triggered by the largest DSFs and they obscure the peaks of suprathreshold DSFs, we conservatively estimated the amplitudes of all suprathreshold DSFs to be equal to the largest subthreshold DSF measured across the entire 30-40 second recording period. Note in both panels that APs and DSFs show little apparent difference from those recorded under current clamp (see preceding figures). (B) Significant DSF amplitudes in SA neurons compared with No SA neurons, both for smaller (left panel, n = 49 versus 203, respectively) and larger DSFs (right panel, n = 47 versus 182, respectively). (C) Significant enhancement of the frequency of larger DSFs in SA neurons versus No SA neurons (n = 48 versus 202, respectively, left and right panels). Red lines indicate group medians, blue lines interquartile range. Each point represents the mean value for all DSFs recorded from 1 neuron across 30-40 seconds. Each group comparison was made using a Mann Whitney U test. DSF, depolarizing spontaneous fluctuation; MP, membrane potential; SA, spontaneous activity.

### 3.4. DSF generation depends upon extracellular Na^+^ and is influenced by extracellular Ca^2+^

Our focus in this study is on the transient depolarizing phase of DSFs, which is important for driving membrane potential to AP threshold in at least some spontaneous pain conditions (see Discussion). In principle, depolarization can be produced by decreasing resting outward current and/or by increasing inward current. Outward current underlying nociceptor RMP is largely conducted by various K^+^ channels ^18^. If DSF generation depended upon spontaneous closing of K^+^ channels, then a large experimental reduction in K^+^ current should occlude any depolarizing effects of spontaneous channel closing, greatly reducing consequent DSFs. However, when we substantially reduced K^+^ current by replacing KCl in the patch pipette with CsCl, DSFs were still observed when held under current clamp at ~-45 mV (unpublished observations). This indicated that the depolarizing phase of DSF generation does not require transient reduction of K^+^ current. These data are part of a study of the contributions of K^+^ channel function to DSF properties that will be reported elsewhere.

We tested the likely importance of extracellular Na^+^ and/or Ca^2+^ for DSF generation by sequentially perfusing DRG neurons with normal extracellular solution (ECS), followed by solutions deficient in extracellular Na^+^ or Ca^2+^, then by a solution largely lacking both cations, and finally by repeating perfusion with the first two solutions. Membrane potential was held at −50 mV by low frequency voltage-clamp. Following perfusion with normal ECS (perfusion 1), the neurons were perfused with a low-Na^+^ ECS (perfusion 2) that contained only 2.2 mM Na^+^ (with the remainder substituted by choline^+^), bringing the Na^+^ reversal potential close to the holding potential, thus greatly decreasing the driving force on Na^+^. The calculated reversal potential was approximately −45 mV, assuming the intracellular concentration was equal to the pipette concentration of 13.8 mM Na^+^. Compared to ECS (perfusion 1), the mean amplitudes of smaller DSFs (1.5-3 mV) were reduced significantly in low-Na^+^ ECS (perfusion 2, **Fig. 6A, B**). In contrast, the mean amplitudes of larger DSFs (>3 mV) showed little or no change in low-Na^+^ ECS (**Fig. 6A, C**). The largest effect of low-Na^+^ ECS was on the frequencies of smaller DSFs, which decreased in all tested neurons (**Fig. 6A, D**). However, the frequencies of larger DSFs showed no significant change (**Fig. 6A, E**), indicating that extracellular Na^+^ contributes less to the larger DSFs than to the smaller DSFs. When the neurons were then exposed to a solution (perfusion 3) low in both extracellular Na^+^ and Ca^2+^ (2.2 mM Na^+^ and ~0 mM Ca^2+^ plus 1 mM EGTA), all the larger DSFs were eliminated along with the vast majority of smaller DSFs (**Fig. 6A, C, D, E**). Subsequent perfusion with low-Na^+^ ECS (perfusion 4) and then with ECS (perfusion 5) reinstated both smaller and larger DSFs, and the larger DSFs tended to rebound with higher mean frequencies and larger mean amplitudes (**Fig. 6A-E**). Small spikes usually appeared during the rebound in the low-Na^+^ ECS, and full APs appeared during perfusion 5 with ECS (**Fig. 6A**). Importantly, these results show a dependence of the smaller DSFs (1.5-3.0 mV, representing a majority of all DSFs analyzed) on Na^+^ influx. The DSFs and small spikes present in our low-Na^+^ solutions might be generated within neuronal subpopulations expressing channels having significant permeability to Ca^2+^ and/or to choline^+^.

**Figure 6.**
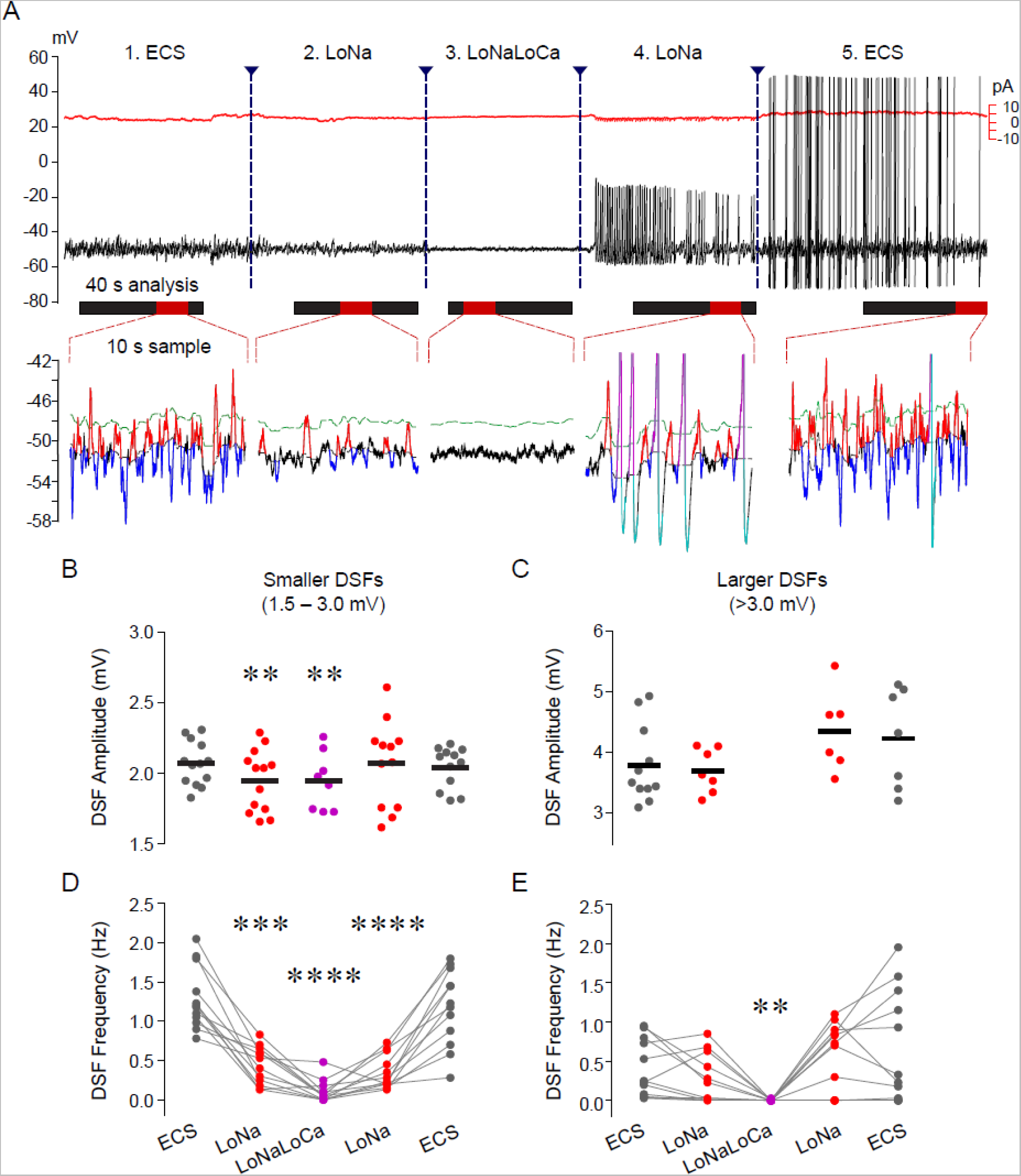
DSF generation depends upon extracellular Na^+^ and Ca^2+^. (A) Example of a DRG neuron perfused sequentially with solutions containing 1) ECS, 2) low-Na^+^ ECS (choline-Cl substituting for all but 2.2 mM of NaCl), 3) low-Na^+^/low-Ca^2+^ ECS (choline-Cl with 0 mM Ca^2+^ plus 1 mM EGTA), 4) low-Na^+^ ECS, and 5) ECS. Representative spontaneous fluctuations during 10-second samples (red bars) within each 40-second analysis window (black bars) are shown below the original recording. (B) Mean DSF amplitudes during each analysis window across perfusions. Not all neurons showed detectable DSFs during each perfusion, preventing repeated measures tests across all perfusions, so planned comparisons were made using paired t-tests between responses to ECS (n = 13) and responses to either low-Na^+^ ECS (n = 13) or to low-Na^+^/low-Ca^2+^ ECS (n = 8). **, *P* < 0.005, after Bonferroni correction. (C) Lack of significant effects on the amplitudes of larger DSFs using the same analyses as in panel B (paired t-tests, n= 11, 7, and 0 in the first 3 perfusions). Larger DSFs were eliminated by low-Na^+^/low-Ca^2+^ ECS. (D) Changes in mean frequencies for smaller DSFs during each analysis window across perfusions (n = 12 for each perfusion). (E) Changes in mean frequencies for larger DSFs during each analysis window across perfusions (n = 12 for each perfusion). For panels D and E, significant differences compared to the first ECS perfusion were assessed with 1-way ANOVA for repeated measures followed by Dunnett’s post-tests. **, *P* < 0.01; ***, *P* < 0.001, ****, *P* < 0.0001. ECS, extracellular solution; LoNa, low-Na^+^ ECS; LoNaLoCa, low-Na^+^/low-Ca^2+^ ECS; DSF, depolarizing spontaneous fluctuation.

Somewhat different results were found when the perfusion sequence included low-Ca^2+^ ECS instead of low-Na^+^ ECS (**Fig. 7A**). Following ECS (perfusion 1), low-Ca^2+^ ECS (perfusion 2) had no significant effect on the amplitudes or frequencies of smaller or larger DSFs (**Fig. 7B, C**). Once again, however, perfusion 3 with an ECS low in both Na^+^ and Ca^2+^ reversibly eliminated all the large DSFs along with the vast majority of smaller DSFs (**Fig. 7A, B, D, E**). Reperfusion with low-Ca^2+^ ECS (perfusion 4) and then with ECS (perfusion 5) reinstated both smaller and larger DSFs, without evidence of rebound enhancement for smaller DSFs but significant rebound enhancement of larger DSFs during the second ECS exposure (perfusion 5, **Fig.7B-E**). These observations show that Ca^2+^ influx is not necessary for DSF generation when extracellular Na^+^ is available, but it may be essential in the absence of an effective driving force for Na^+^ influx. Together with the results shown in **Figure 6**, these findings suggest that, under basal conditions, DSFs are largely generated by spontaneous opening of channels permeable to Na^+^ and channels permeable to both Na^+^ and Ca^2+^. In addition, or alternatively, direct or indirect interactions between populations of DSF-driving channels separately permeable to either Na^+^ or Ca^2+^ might account for the much stronger effects of concurrently lowering Na^+^ and Ca^2+^ in the ECS compared with lowering either cation alone.

**Figure 7.**
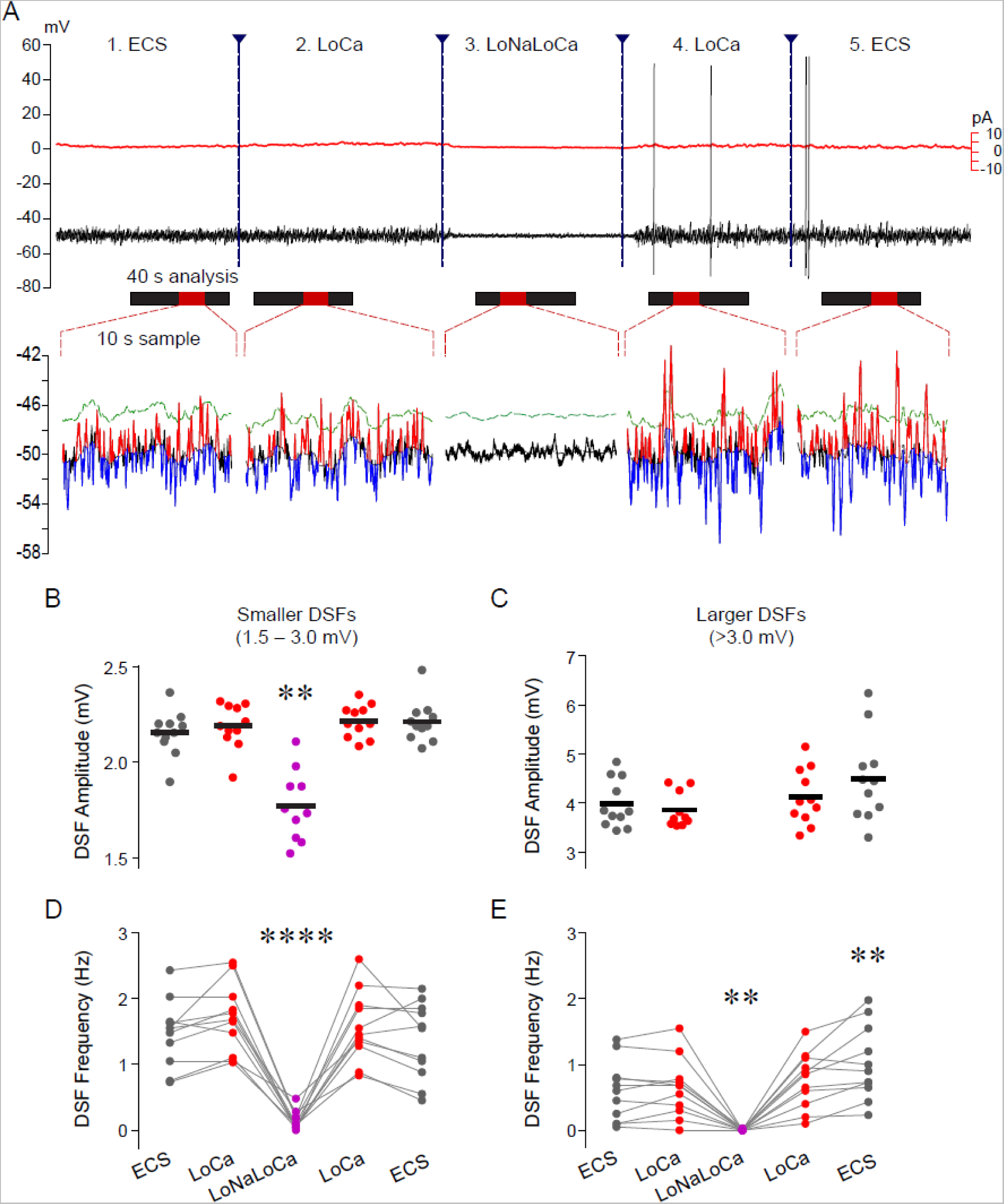
DSF generation depends upon extracellular Na^+^ and Ca^2+^ as shown by the effects of substituting extracellular Ca^2+^ followed by substituting both Ca^2+^ and Na^+^. (A) Example of a DRG neuron perfused sequentially with solutions containing 1) ECS, 2) low-Ca^2+^ ECS (0 mM Ca^2+^ plus 1 mM EGTA), 3) low-Na^+^/low-Ca^2+^ ECS, 4) low-Ca^2+^ ECS, and 5) ECS. Representative spontaneous fluctuations during 10-second samples within each 40-second analysis window are shown below the original recording. (B) Mean DSF amplitudes during each analysis window across perfusions, showing significant reduction in low-Na^+^/low-Ca^2+^. Planned comparisons were made using paired t-tests between responses to ECS (n = 11) and responses to either low-Na^+^ ECS (n = 11) or to low-Na^+^/low-Ca^2+^ ECS (n = 10). **, *P* < 0.005, after Bonferroni correction. (C) Lack of significant effects on the amplitudes of larger DSFs, except for the complete elimination of larger DSFs by low-Na^+^/low-Ca^2+^ ECS (n = 11, 10, 0, 11, 11 in successive perfusions). (D) Changes in mean frequencies for smaller DSFs during each analysis window across perfusions (n = 11). (E) Changes in mean frequencies for larger DSFs during each analysis window across perfusions (n = 11). For panels D and E, significant differences compared to the first ECS perfusion were assessed with 1-way ANOVA for repeated measures followed by Dunnett’s post-tests. **, *P* < 0.01, ****, *P* < 0.0001. ECS, extracellular solution; LoCa, low-Ca^2+^ ECS; LoNaLoCa, low-Na^+^/low-Ca^2+^ ECS; DSF, depolarizing spontaneous fluctuation.

### 3.5. DSF generation in capsaicin-sensitive nociceptors depends upon multiple types of channels permeable to Na^+^ and/or Ca^2+^

Our ion substitution experiments showed that DSFs depend upon the influx of both Na^+^ and Ca^2+^. To screen numerous channel types known to be permeable to Na^+^ and/or Ca^2+^ and to be expressed in primary nociceptors, we primarily employed a pharmacological approach, using concentrations of inhibitors previously reported to reduce excitability or inward currents in nociceptors, and we focused on neurons showing clear depolarizing responses to perfusion of 1 μM capsaicin (probable peptidergic nociceptors in mice ^13, 22, 25^) delivered at the end of the recording. After at least 75 seconds of perfusion with ECS, a selected ion channel inhibitor was perfused for at least 3 minutes. DSF properties were measured under low-frequency voltage clamp at −45 or −50 mV for 40 seconds beginning at least 30 seconds after starting perfusion of ECS and at least 150 seconds after starting perfusion of the inhibitor so that any steady-state effects of inhibitor exposure could be observed.

We first sought evidence for potential contributions to DSFs of the three VGSCs, Nav1.7, Nav1.8, and Nav1.9, that have the most prominent roles in nociceptor function ^17^. We began with Nav1.8 because of our earlier finding that antisense knockdown of Nav1.8 largely eliminated nociceptor SA after SCI ^72^. **Figure 8A** shows examples of responses during ECS perfusion and then after switching to either ECS again (control) or 1 μM A-803467, a specific inhibitor of Nav1.8 channels ^33^. In these examples there was no change in mean DSF amplitude or frequency during the control ECS perfusion. In contrast, while the illustrated neuron exhibited a 1.9% increase in mean amplitude of the smaller DSFs (1.5-3 mV) when perfused with A-803467, it showed a 15.8% decrease in amplitude of the larger DSFs (>3 mV), along with a 5.8% decrease in the frequency of smaller DSFs and a 35.6% decrease in larger DSF frequency. Summary data for all neurons tested (**Tables 1** and **2**) show that control perfusion with ECS had no significant effect on the mean amplitudes or frequencies of either smaller or larger DSFs, while A-803467 decreased the mean amplitude and frequency of larger DSFs by 9.8% and 27.7%, respectively, without affecting smaller DSFs. Similarly, the Nav1.7 inhibitor, ProTx II ^19, 56^, significantly decreased the amplitude and frequency of larger DSFs by 11.2% and 25.8%, respectively. Effective and selective inhibitors of Nav1.9 are not available, so we compared DSF properties under low-frequency voltage clamp of ECS-perfused neurons from wild-type mice and mice with global knockout of Nav1.9 (confirmed by genotyping). Interestingly, in neurons from Nav1.9^−/−^ mice, smaller DSFs exhibited slightly but significantly larger amplitudes (3.2%) but a 31.2% lower frequency than those in neurons from wild-type mice. No significant differences were found in larger DSFs (**Tables 1** and **2**).

**Figure 8.**
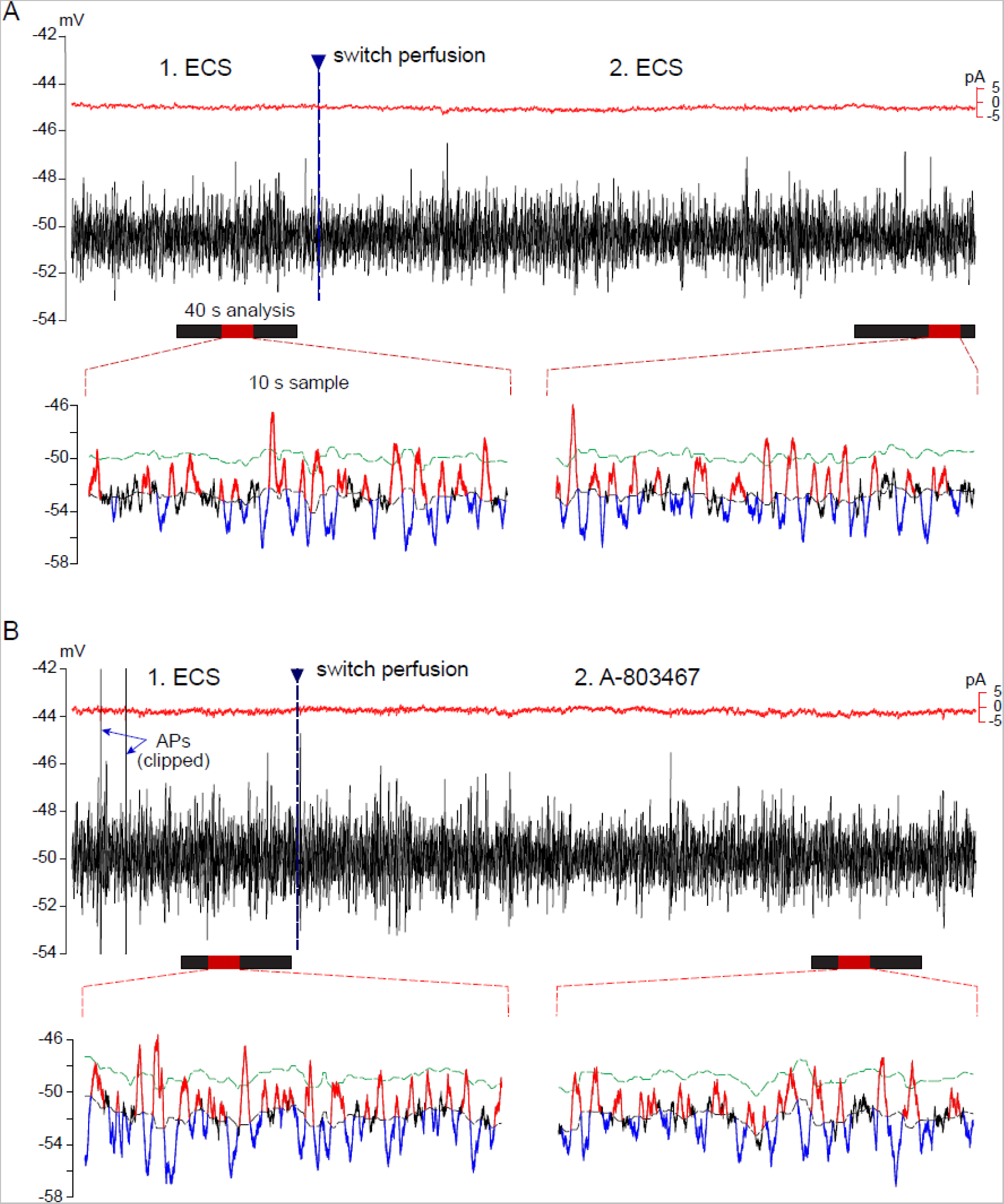
DSFs can be reduced by perfusion with VGSC blockers: example of modest reduction by a Nav1.8 blocker, A-803467. (A) ECS control. Analysis (40-second windows, black bars, containing 10-second examples, red bars) was performed shortly before switching perfusion and then beginning 3 minutes afterwards. (B) Small reduction of DSF amplitude and the frequency of larger DSFs measured 3 minutes after switching to A-803467 (1 μM). APs, action potentials; ECS, extracellular solution.

**Table 1:** Effects of ion channel blockers on smaller DSFs (1.5 – 3 mV)

| Target (blocker) | Amplitudes (mV) |  |  |  | Frequencies (Hz) |  |  |  |  |
| --- | --- | --- | --- | --- | --- | --- | --- | --- | --- |
|  | ECS | Treatment | Change % | P | ECS | Treatment | Change % | P | n |
| Control (ECS repeated) | 2.1 | 2.1 | 1.1 | 0.167 | 1.6 | 1.5 | -6.5 | 0.175 | 31 |
| Nav1.7 (20 nM ProTX II) | 2.1 | 2.1 | 0.0 | 0.980 | 1.6 | 1.6 | -5.5 | 0.119 | 17 |
| Nav1.8 (1 μM A-803467) | 2.1 | 2.2 | 0.6 | 0.571 | 1.5 | 1.5 | -2.8 | 0.617 | 15 |
| Nav1.9 (Nav1.9<sup>-/-</sup>) | 2.0 WT | 2.1 KO | 3.2 | <0.01 | 2.6 WT | 1.8 KO | -31.2 | <0.001 | 42; 45 |
| HCN (ZD-7288) | 1.9 | 2.0 | 1.7 | 0.090 | 1.3 | 1.2 | -7.3 | 0.214 | 20 |
| NALCN (10-50 μM, GdCl<sub>3</sub>) | 2.1 | 2.0 | -1.9 | 0.507 | 1.1 | 0.8 | -24.5 | <0.01 | 8 |
| NALCN (10 μM L-703606) | 2.1 | 2.0 | -6.1 | 0.081 | 1.5 | 0.9 | -36.6 | <0.01 | 7 |
| TRPV1 (10 μM AMG-9810) | 2.1 | 2.2 | 0.4 | 0.677 | 1.7 | 1.3 | -26.0 | <0.01 | 31 |
| TRPA1 (10 μM A-967079) | 2.0 | 2.0 | 0.7 | 0.691 | 1.2 | 0.8 | -31.6 | <0.001 | 19 |
| TRPM3 (1 μM ononetin) | 2.2 | 2.2 | 0.8 | 0.398 | 1.5 | 1.4 | -6.8 | 0.363 | 23 |
| TRPM4 (50 μM CBA) | 2.1 | 2.1 | 0.4 | 0.722 | 1.4 | 1.5 | 12.1 | 0.351 | 20 |
| TRPC4/5 (0.1 μM pico-145) | 2.1 | 2.1 | -0.2 | 0.836 | 1.6 | 1.4 | -15.3 | 0.126 | 22 |
| Cav-T, (1 μM TTA-P2) | 2.1 | 2.1 | -0.7 | 0.586 | 1.5 | 1.4 | -7.0 | 0.168 | 19 |
| Cav -L, (10 μM nimodipine) | 2.1 | 2.1 | 0.4 | 0.823 | 2.2 | 2.3 | 2.3 | 0.882 | 7 |
| Cav -N, (0.5 μM ω-conotoxin GVIA) | 2.1 | 2.1 | -1.9 | 0.095 | 1.7 | 1.7 | 0.2 | 0.981 | 13 |

HCN channels are permeable to Na^+^ and K^+^, and HCN channels in nociceptors have been linked to inflammatory and neuropathic pain ^21, 41^. Moreover, they contribute to input resistance in small DRG neurons of mice ^15^. However, we found that the selective HCN blocker ZD-7288 ^10^, had no significant effects on the amplitude or frequency of either smaller or larger DSFs (**Tables 1** and **2**). Mammalian nociceptors also express numerous nonselective cation channels that are permeable to both Na^+^ and Ca^2+^ as well as K^+^. Of special interest is the Na^+^ leak channel NALCN, a nonselective monovalent cation channel with a relatively high probability of being open at RMP that has been implicated in driving pain ^20, 43, 74^. Highly specific NALCN inhibitors are not available, so we examined the effects of an effective but nonselective NALCN blocker, Gd^3+^ ^45^, and a somewhat more selective NALCN inhibitor, L-703606 ^28^. Gd^3+^ decreased the amplitude and frequency of larger DSFs by 12.3% and 64.6% (**Table 2**). Gd^3+^ also reduced the frequency of smaller DSFs by 24.5%. Similarly, L-703606 decreased the amplitude and frequency of larger DSFs by 12.2% and 54.8%, respectively, and it reduced the frequency of smaller DSFs by 36.6% (**Tables 1 and 2**). The large magnitudes of these reductions are consistent with each of these nonspecific inhibitors reducing the activity of multiple channel types in addition to NALCN.

**Table 2:** Effects of ion channel blockers on larger DSFs (> 3 mV)

| Target (blocker) | Amplitudes (mV) |  |  |  | Frequencies (Hz) |  |  |  |  |
| --- | --- | --- | --- | --- | --- | --- | --- | --- | --- |
|  | ECS | Treatment | Change % | P | ECS | Treatment | Change % | P | n |
| Control (ECS repeated) | 4.0 | 4.0 | 0.9 | 0.770 | 0.6 | 0.6 | 3.5 | 0.767 | 30 |
| Nav1.7 (20 nM ProTX II) | 4.2 | 3.7 | -11.2 | <0.01 | 0.5 | 0.4 | -25.8 | 0.050 | 14 |
| Nav1.8 (1 μM A-803467) | 4.5 | 4.0 | -9.8 | <0.01 | 1.1 | 0.8 | -27.7 | <0.01 | 15 |
| Nav1.9 (Nav1.9<sup>-/-</sup>) | 3.9 WT | 3.9 KO | 0.3 | 0.931 | 0.4 WT | 0.5 KO | 26.1 | 0.222 | 41; 45 |
| HCN (ZD-7288) | 3.6 | 3.7 | 1.4 | 0.745 | 0.3 | 0.4 | 18.4 | 0.483 | 17 |
| NALCN (10-50 μM, GdCl<sub>3</sub>) | 4.3 | 3.8 | -12.3 | <0.05 | 0.5 | 0.2 | -64.6 | <0.05 | 8 |
| NALCN (10 μM L-703606) | 4.2 | 3.7 | -12.2 | <0.05 | 0.6 | 0.3 | -54.8 | 0.069 | 5 |
| TRPV1 (10 μM AMG-9810) | 4.4 | 4.1 | -6.4 | <0.01 | 0.9 | 0.7 | -24.7 | <0.05 | 30 |
| TRPA1 (10 μM A-967079) | 3.9 | 3.9 | -0.1 | 0.989 | 0.3 | 0.2 | -39.8 | 0.091 | 19 |
| TRPM3 (1 μM ononetin) | 4.3 | 4.1 | -4.0 | 0.166 | 0.8 | 0.7 | -13.6 | 0.187 | 22 |
| TRPM4 (50 μM CBA) | 4.1 | 3.9 | -5.7 | <0.01 | 0.6 | 0.4 | -23.4 | 0.073 | 20 |
| TRPC4/5 (0.1 μM pico-145) | 4.1 | 4.2 | 2.1 | 0.458 | 0.8 | 0.7 | -8.4 | 0.484 | 22 |
| Cav-T, (1 μM TTA-P2) | 3.8 | 3.8 | 1.1 | 0.795 | 0.4 | 0.3 | -20.0 | 0.300 | 18 |
| Cav -L, (10 μM nimodipine) | 4.2 | 3.9 | -5.8 | 0.214 | 1.0 | 1.1 | 9.0 | 0.505 | 7 |
| Cav -N, (0.5 μM ω-conotoxin GVIA) | 4.2 | 4.0 | -6.2 | <0.05 | 0.9 | 0.6 | -26.8 | <0.01 | 13 |

Nonselective cation channels that are represented in nociceptors also include the transient receptor potential (TRP) family, at least three members of which are known to be important for nociception and pain: TRPV1, TRPA1, and TRPM3 ^4^. A specific TRPV1 inhibitor, AMG-9810 ^27^, decreased the amplitude and frequency of larger DSFs by 6.4% and 24.7%, respectively (**Table 2**). AMG-9810 also reduced the frequency of smaller DSFs by 26.0%. A specific TRPA1 inhibitor, A-967079 ^47^, had no significant effects on the amplitude of smaller or larger DSFs, but reduced the frequency of smaller DSFs by 31.6% and exhibited a nonsignificant trend to reduce the frequency of larger DSFs (**Tables 1 and 2**). A specific TRPM3 inhibitor, ononetin ^60^, had no significant effects on the amplitude or frequency of smaller or larger DSFs (**Tables 1 and 2**). A specific TRPM4 inhibitor, CBA ^53^, reduced the amplitude of larger DSFs by 5.7% without having significant effects on their frequency, or on the amplitude or frequency of smaller DSFs (**Tables 1 and 2**). Finally, an inhibitor specific to TRPC4 and TRPC5, Pico-145, had no significant effects on the amplitude or frequency of smaller or larger DSFs (**Tables 1 and 2**).

In addition, we tested inhibitors of VGCCs that underlie T-type, L-type, and N-type Ca^2+^ currents having known roles in nociceptor function and pain ^30, 35^. Neither an inhibitor of T-type current, TTA-P2 ^14^, nor of L-type current, nimodipine ^57^, had significant effects on the amplitude or frequency of either smaller or larger DSFs (**Tables 1 and 2**). However, a relatively specific inhibitor of N-type current, ω-conotoxin GVIA ^37, 46^, decreased the amplitude and frequency of larger DSFs by 6.2% and 26.8%, respectively (**Table 2**).

Together, these results indicate that the depolarizing phase of DSFs occurring at holding potentials of −45 to −50 mV in capsaicin-sensitive nociceptors isolated from previously uninjured mice involves a diverse set of ion channels permeable to Na^+^ and/or Ca^2+^. Although conclusive evidence about the relative contributions to DSFs from specific channels will require more selective tests than used in this broad pharmacological screen, we can conclude that DSF generation at membrane potentials between RMP and AP threshold in nociceptors depends upon spontaneous fluctuations of inward current through multiple types of voltage-gated and nonselective cation channels.

## 4. Discussion

This study shows that, across a wide range of membrane potentials, capsaicin-sensitive nociceptors exhibit both high input resistance and DSFs that require background activity of diverse ion channels permeable to Na^+^ and/or Ca^2+^. These specializations explain how electrically silent nociceptors are poised to develop ongoing electrical activity that can promote spontaneous pain.

### 4.1. Nociceptor somata exhibit unusually high input resistance

Because the amplitudes of fluctuations in membrane potential occurring spontaneously in nociceptor somata are determined by changes in the product of the net current flowing across the plasma membrane and the neuron’s input resistance, higher input resistance will increase the amplitude of any DSFs caused by spontaneous, transient opening of somal membrane ion channels. We found in mouse nociceptor somata that input resistance measured at holding potentials of approximately −50, −60, or −70 mV was often ≥ 3 GΩ when measured with hyperpolarizing pulses, and usually 1 to 2 GΩ when measured with depolarizing pulses reaching a level (~ −40 mV) where large DSFs can approach AP threshold ^9^. These resistance values are quite high compared with most reported values from whole-cell patch recordings for neurons with similar soma size (10-25 μm diameter). For example, hippocampal, cortical, cerebellar, brainstem, and spinal neurons recorded in slices or in culture usually exhibit input resistances at least 10-fold lower ^11, 12, 31, 38, 73^. Part of this difference reflects the background synaptic conductance in central neurons recorded in slices, or in vivo where input resistance is even lower ^24^, whereas DRG neurons don’t receive synapses in vivo or in vitro. Lacking dendrites, a DRG neuron soma is in significant electrical continuity only with its stem axon connected to the T-junction and its peripheral and central axonal branches. In addition, dissociation of DRG neurons may eliminate neuromodulatory inputs that might reduce input resistance.

Compared with most DRG neurons, the mean values previously reported for somal input resistance tested with hyperpolarizing pulses under whole-cell current clamp are higher in mouse DRG neurons, which range from approximately 350 to 750 MΩ ^15, 23, 34, 36, 54^, which are lower than those reported here. This discrepancy probably reflects methodological differences, such as the composition of the solutions in the recording pipette and ECS, or the period after dissociation when the recordings were made ^52^. In most studies, recording occurred on the same day as dissociation ^23, 34, 36, 54^, whereas we waited 1 day. Another difference may be in neuronal selection criteria. Input resistance is negatively correlated with input capacitance, and we selected for relatively small DRG neurons with median input capacitance of 16 pF. Input resistance is also directly affected by the seal resistance between the membrane and pipette. We required a seal resistance ≥ 3 GΩ, whereas many investigators use a 1 GΩ minimum or fail to report any minimum. The unusually high input resistances we found in probable nociceptors under our dissociation conditions may be somewhat higher than those occurring in vivo because of the loss of axons, a lack of background neuromodulation, or cellular stress caused by isolation and recording procedures ^1, 52, 75^. Nonetheless, anatomical features of DRG neurons, the relatively high DRG neuron input resistances reported by others, and the even higher input resistances we found suggest that high somal input resistance is a functional specialization in nociceptors that enhances somal DSF generation.

### 4.2. Diverse channel types contribute to DSF generation in nociceptor somata

Our ion substitution results on neurons from previously uninjured mice suggest that DSFs are mainly generated by channels permeable to Na^+^ and/or to both Na^+^ and Ca^2+^. An earlier study of rat nociceptors indicated that, in addition to prominent outward current through several K^+^ channel families, RMP is set by background inward current through HCN channels, Nav1.8, and TTX-sensitive Na^+^ channels ^18^. All these Na^+^-permeable channels are potential contributors to the depolarizing phase of DSFs at RMP. However, consistent with the hyperpolarized voltage sensitivity of HCN channels ^21^, our results suggest little or no contribution of HCN channels to DSF generation at holding potentials (−50 to −45 mV) where larger DSFs can approach AP threshold. In contrast, significant decreases in the amplitude and/or frequency of DSFs produced by relatively specific inhibitors of Nav1.7, Nav1.8, TRPV1, TRPA1, TRPM4, and N-type Ca^2+^ channels suggest that these channels contribute to DSF generation under our conditions. Furthermore, contributions of NALCN channels are suggested by decreases in DSF amplitude and frequency from effective but nonspecific inhibitors of these channels. Interestingly, global knockout of Nav1.9 channels significantly reduced the frequency of smaller but not larger DSFs compared to those in wild-type neurons, whereas inhibitors of Nav1.7, Nav1.8, and N-type Ca^2+^ channels only reduced larger DSFs. Thus, Nav1.9 (which has a relatively hyperpolarized activation range) may contribute to the less depolarized phase of DSFs while Nav1.7, Nav1.8, and N-type Ca^2+^ channels having more depolarized activation ranges may contribute to the more depolarized phases of larger DSFs. TRPV1 appears to contribute both to smaller and larger DSFs, TRPA1 to smaller DSFs, and TRPM4 to larger DSFs.

For most channel types, the applied inhibitor had a stronger effect on the frequency than amplitude of DSFs. The low frequencies of DSFs might reflect the necessity of multiple channel types to open spontaneously and concurrently to generate DSFs exceeding detection thresholds (1.5 mV for smaller DSFs, 3 mV for larger DSFs). During pharmacological inhibition, a small, further decrease in the already low probability of opening for any channel type might substantially increase the average time between sufficient concurrent openings of multiple channels to produce a DSF exceeding the set threshold. While some nonspecific effects are likely with all drugs (especially the available NALCN inhibitors), we can conclude that DSF generation in previously uninjured nociceptors at membrane potentials between RMP and AP threshold depends upon spontaneous fluctuations of inward current through multiple types of voltage-gated and nonselective cation channels known to be permeable to Na^+^, to Ca^2+^, and/or to both Na^+^ and Ca^2+^.

### 4.3. Background activity of multiple excitatory ion channels coupled with high input resistance readies nociceptor somata for ongoing discharge

In mammalian neurons, the soma is usually less excitable than the axonal initial segment, or is even inexcitable ^55^, as is the case in most invertebrates other than gastropod molluscs. In nociceptors, however, APs can be generated in the soma, which in principle will easily propagate from there into a nociceptor’s central axon ^62^. While passive electrical properties of DRG neuron somata may serve to filter afferent APs ^29^, this possible function does not require somal AP generation. Instead, persistent activity generated in vivo by DRG neuronal somata is associated with neuropathic conditions ^65, 68^, including amputation in humans ^64^ and SCI in rats ^7^. Spontaneous activity induced in vivo and observed later in vitro has been linked to increased input resistance in rat nociceptors ^7^ and to enhanced DSFs in nociceptors from rodents ^6, 9, 26, 42, 51^ and humans ^50^.

Nociceptor somata are normally electrically silent except when activated by axonal Aps evoked by peripheral stimulation. However, the present findings indicate that nociceptor somata may be continuously poised to develop low-frequency, irregular, ongoing activity under conditions such as exposure to inflammatory signals ^44, 51, 70^ or neuropathy ^65, 68^. Their high input resistance enables relatively small current fluctuations to produce significant DSFs, and the DSFs will be enhanced further with any additional increases in input resistance and/or spontaneous inward current fluctuations. Decreases in nociceptor K^+^ conductances that would both increase input resistance and depolarize RMP closer to AP threshold have been found in numerous neuropathic conditions ^59^, as have increases in the activity of voltage-gated Na^+^ and Ca^2+^ channels and of TRP channels ^8, 32, 58^ that might contribute to DSFs, especially larger DSFs that can approach AP threshold. Our present finding that SA under basal conditions is associated with increased input resistance, depolarized RMP, and enhanced DSFs compared with neurons without SA (see also ^7, 9, 50, 51^) supports the hypothesis that ongoing activity in nociceptors can develop in painful conditions because of further increases in somal input resistance and in spontaneous inward currents through one or more cation channels permeable to Na^+^, to Ca^2+^, and/or to both Na^+^ and Ca^2+^. Because diverse channel types have sufficient combined background activity to generate DSFs stochastically that sometimes approach AP threshold under normal conditions, small increases in input resistance and/or increases in the activity of inward current channels might easily transform an electrically silent soma into one generating low-frequency, irregular discharge. This unusual capability of nociceptor somata might have been selected during evolution as a mechanism for low-frequency ongoing activity in numerous nociceptors to inform the brain of long-lasting vulnerability of an individual following injury severe enough to disconnect peripheral terminals, and this somal state might also be induced in other neuropathic and inflammatory conditions ^66, 67, 69^. Characterization of the contributions of the ion channel types implicated in the present study (and other channels) to the generation of large DSFs under pathological conditions may point to molecular targets offering promise for more effective treatment of persistent ongoing pain.

## Acknowledgements

We thank Dr. Murat Gorgun and Kayla Johnson for expert technical assistance.

